# Insufficient Eye Tracking Data Leads to Errors in Evaluating Typical and Atypical Fixation Preferences

**DOI:** 10.1101/2020.09.21.306621

**Authors:** Gabrielle E. Reimann, Catherine Walsh, Kelsey D. Csumitta, Patrick McClure, Francisco Pereira, Alex Martin, Michal Ramot

## Abstract

Eye tracking provides insights into social processing and its deficits in disorders such as autism spectrum disorder (ASD), especially in conjunction with dynamic, naturalistic stimuli. However, reliance on manual stimuli segmentation severely limits scalability. We assessed how the amount of available data impacts individual reliability of fixation preference for different facial features, and the effect of this reliability on between-group differences. We trained an artificial neural network to segment 22 Hollywood movie clips (7410 frames). We then analyzed fixation preferences in typically developing participants and participants with ASD as we incrementally introduced movie data for analysis. Although fixations were initially variable, results stabilized as more data was added. Additionally, while those with ASD displayed significantly fewer face-centered fixations (*p*<.001), they did not differ in eye or mouth fixations. Our results highlight the validity of treating fixation preferences as a stable individual trait, and the risk of misinterpretation with insufficient data.

---

Eye movement data plays an integral role in understanding social processing deficits in neurodevelopmental disorders, including autism spectrum disorder (ASD). These data commonly indicate that individuals with ASD do not view social interactions or process facial features the same way as their typically developing (TD) peers. However, the specifics of eye gaze differences in ASD have been markedly inconsistent. For example, some studies report reduced eye region fixation time in ASD^1,2^. whereas others report no differences between TD individuals and those with ASD^3,4^. Mouth fixations findings are inconsistent as well. Some studies report that individuals with ASD show increased mouth-looking, and even suggest increased mouth fixations serve to compensate for communication deficits deriving from reduced eye region fixation^5^. However, other studies report decreased attention to or no differences in mouth fixation times between TD individuals and those with ASD, regardless of whether stimuli are dynamic or static, or whether they depict multiple or single-persons^6,7,8^.

These inconsistent findings may be in part due to differences in eye tracking methods, particularly techniques for stimuli segmentation. Most researchers rely on manual coding to define regions of interest (e.g. face components, background objects), ^7,9^ however methodological challenges arise with this approach. Manual segmentations can expose stimuli to human error, cause replication complications, and prove to be extremely time-consuming. These factors impede our ability to aggregate eye tracking results and create a coherent understanding of social processing in ASD. Considering the hassles of dynamic stimuli manual segmentation, researchers in turn opt to use fewer movie clips, or reuse already-segmented clips from prior studies^6,7^. Variability in eye tracking findings may reflect generalizability issues with using few or rehashed movie clips. This begs the question, how much of this variability might be explained by insufficient data? Moreover, how much data is enough to produce reliable findings?

Researchers across various disciplines have addressed concerns surrounding the amount of data needed for reliable results. For example, some researchers have applied this question to driving behavior and traffic safety^10^ while others sought to reliably assess dyadic conflict behavior^11^. These findings then are used to provide evidence-based improvements for variable estimates and the planning of future studies^12,13^. While eye tracking researchers have called for reliable and reproducible methods^14^, we have yet to assess basic questions on how much data is needed to consistently estimate eye tracking fixations.

Further, findings on the amount of data needed to reliably assess individual fixations would deepen our understanding of gaze as a stable individual trait. Prior studies have demonstrated that eye scan paths vary greatly among individuals when attending to the face^15,16^. Yet fixations are remarkably stable within an individual across tasks and even over long periods of time^17,18,19^. Despite these promising findings, it is difficult to understand the long-term stability of fixations in the context of static images alone. To understand fixation as a stable trait, it is necessary to investigate an individual’s gaze using dynamic stimuli for real world applicability.

In the present study, we trained an artificial neural network (ANN) to segment naturalistic dynamic stimuli of 22 movie clips; this algorithm allows for the expeditious segmentation of large amounts of stimuli. Utilizing this approach to a free-viewing eye tracking paradigm, we investigated the stability and robustness of within-subject, within-group, and between-group analyses when introducing incremental amounts of movie data for study. First, we examine each participant’s viewing of single movie clips and assess the consistency of proportion of fixations to core facial features (eyes, nose and mouth). We then introduce the effect of varying movie data amounts to examine how much data is needed to observe within-group stability and reliable between-group differences. If indeed the proportion of time individuals fixate on the different facial features is a stable individual trait, then we hypothesize that results of within-group and between-group analyses will become increasingly consistent and robust as we introduce more data. We expect to see this effect in both TD participants and participants with ASD. Subsequently, we applied the previous analysis’ findings in the context of examining how individuals with ASD view elements of social interactions relative to TD peers, based on a quantity of movie data shown to be sufficient for observing reliable differences.

## Results

### Internal consistency of fixations

First, we sought to analyze how consistent each participant is in his own fixations to the different facial features across 22 movies. To this end, we analyzed individual variation in fixations by calculating the fixation proportion per participant, movie clip, and facial feature, compared to all other participants in their group (see Methods). We then compared these individual fixation proportions across all possible movie pairs (e.g. Movie 1 and Movie 2, Movie 1 and Movie 3). The scatterplots in Figure 3a - c each display an example of how well-correlated the participants are to themselves across two example movies; the correlation coefficient is a measure of within-subject internal consistency across all participants in each group for that particular movie pair. The correlation coefficients for all possible movie pairs are then combined to create the histograms featured in Figure 3a - c. These histograms showcase the individual variability of correlations between all single movie pairs for (TD_median correlation_ = 0.63, ASD_median correlation_ = 0.45), mouth (TD_median correlation_ = 0.59, ASD_median correlation_ = 0.46), and nose (TD_median correlation_ = 0.42, ASD_median correlation_ = 0.28) fixations for ASD and TD groups separately.

**Figure 3a, b, c.**
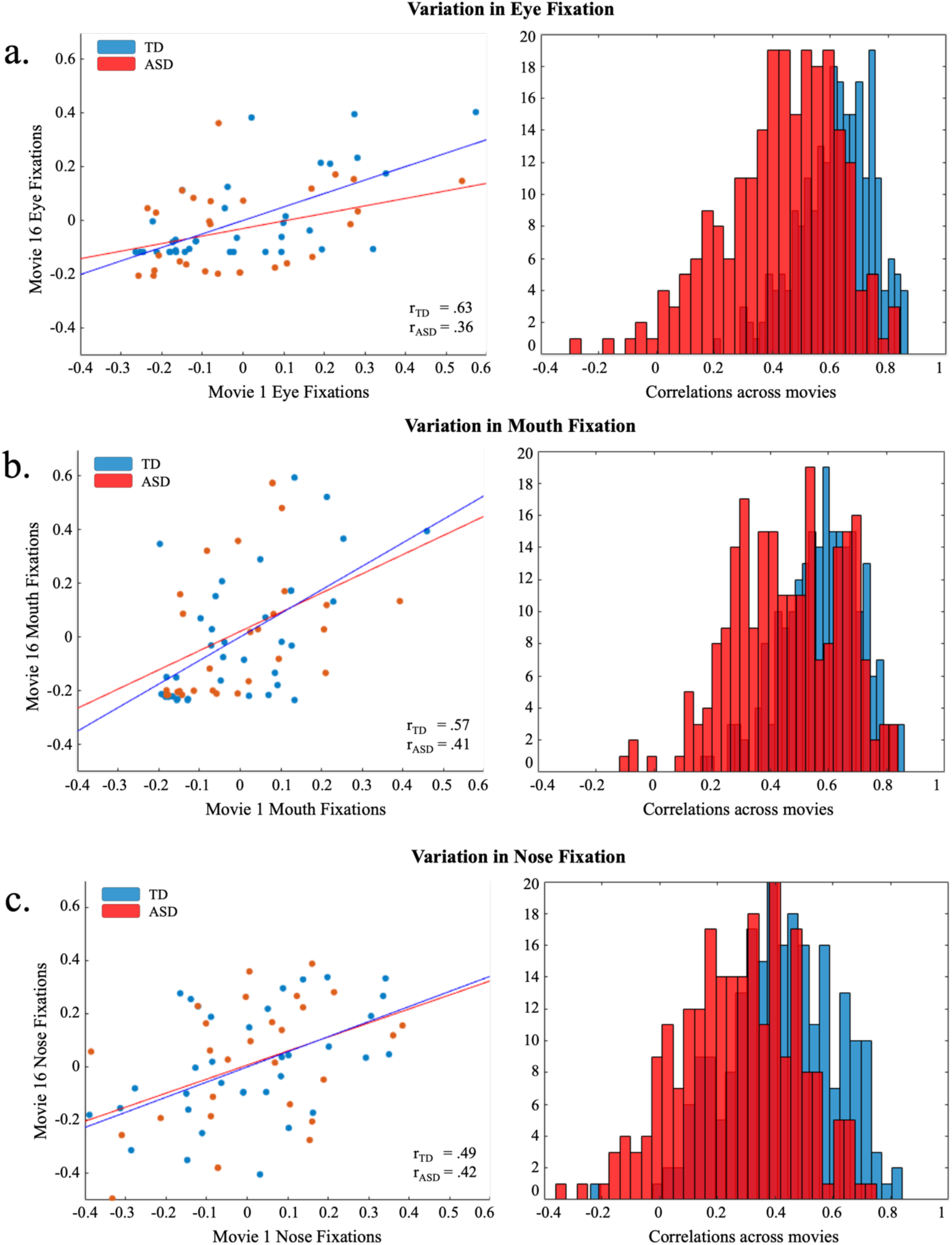
Individual consistency of fixations of TD and ASD participants to the different facial features across 22 movies. The scatterplots display an example of how well-correlated participants are to themselves across two example movies for each of the face labels (Movie 1 versus Movie 16); the correlation coefficient is a measure of within-subject internal consistency across all participants in each group for that particular movie pair. The histograms display the individual variability of correlations between all single movie pairs for eye (Figure 3a.; Td_median_ = 0.63, ASD_median_ = 0.45), mouth (Figure 3b., TD_median_ = 0.59, ASD_median_ = 0.46), and nose (Figure 3c., TD_median_ = 0.42, ASD_median_ = 0.28) fixations for ASD and TD groups separately.

We then carried out a permutation test to assess whether the distributions of these correlation coefficients significantly differ between the TD and ASD groups (see Methods). Compared to TD counterparts, those with ASD showed significantly reduced within-subject internal consistency in facial feature viewing preferences across movie clips (*p* < .001), as well as significantly increased variability (*p* < .001) for eye, nose, and mouth labels. This is in line with previous literature^20,22,23^, which has emphasized inter-subject variations among individuals with ASD. We found a similar result by analyzing the variance of overall fixation time for the different facial features.

### Stability of Within-group Results

To address whether adding more data improves the reliability of fixations and justifies the treatment of gaze allocation to different facial features as a stable trait, we then investigated the consistency of within-group fixations across movies when introducing incremental amounts of movie data to our analyses. We examined this effect on each of the three individual face labels. First, from our 22 movie clips, we randomly selected two sets of three non-overlapping movies combinations, totaling 42 seconds of stimuli. Similar to the previous analysis, we calculated the correlations of the within-group internal consistency of the fixation proportions across these two sets of movies; this was done for TD participants and participants with ASD separately. This process was repeated for 10,000 permutations, with different sets of three movies selected for each permutation. As before, the histograms in Figure 4 display the correlation coefficients for all the different permutations. To assess incremental additions of data, we repeated this process by creating two sets of movies with random combinations of five (70 seconds), eight (112 seconds), and 11 (154 seconds) movies. Figure 4a - c displays the distribution of correlations as the number of movie clips increases for eye, mouth, and nose fixations, respectively. Across each face label, Figure 4 displays the increasing reliability of correlations as data increases.

**Figure 4a, b, c.**
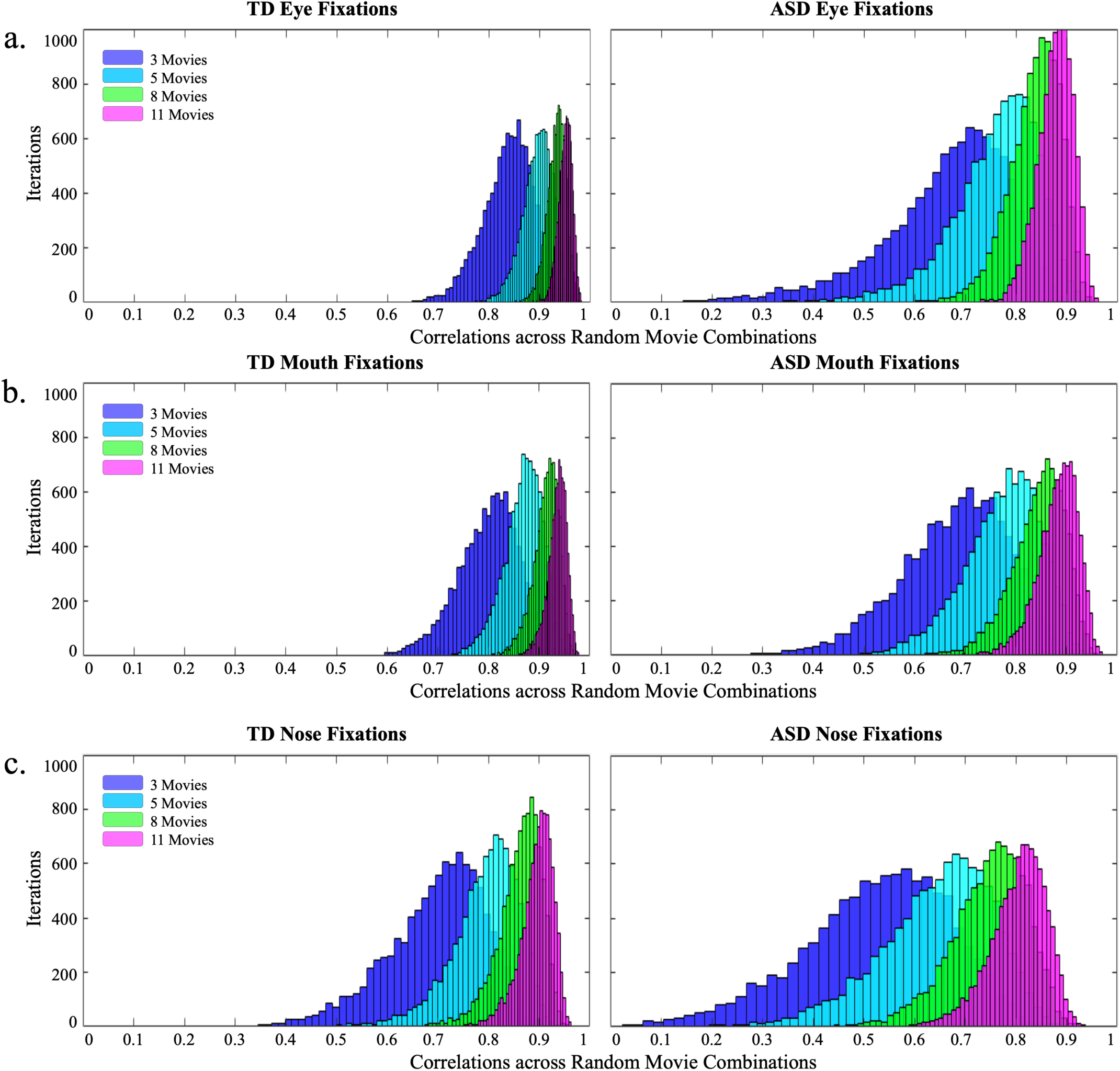
Change in the consistency of within-group fixations across movies when introducing additional movie data for TD (left panel) and ASD (right panel) participants. Histograms reflect the distribution of correlation coefficients of all permutations as the number of movie clips increases for eye (Figure 4a.), mouth (Figure 4b.), and nose (Figure 4c.) fixations, respectively (see methods for details).

Next, we tested whether there is significantly increased reliability when using more data. Using permutation tests, we examined whether the distributions of internal consistency using different numbers of movies (Figure 4) were significantly different from each other (see Methods). Results show that within both the ASD and TD groups, the medians and the variance of the distributions using different numbers of movies were significantly different from each other for each of the face labels (*p*_median_ < 1×10^−4^; *p*_variance_ < 1×10^−4^ for all pairwise comparisons). For a given amount of movie data (i.e. 3 movies, 5 movies etc.), there were also significant differences in the distributions across groups. Those with ASD were significantly less consistent (*p*_median_ < 1×10^−4^) and more variable (*p*_variance_ < 1×10^−4^) than their TD peers across all data amount levels.

### Stability of Between-group Results

Thus far, we have demonstrated that increasing amounts of movie data serves to stabilize individual fixation variation within each group. With this basis, we then examined the effect of this increased stability on the reliability of ASD and TD between-group facial feature fixation differences. First, we examine the variability in between-group differences when using a single movie clip. We used two-sample t-tests to examine between-group differences per face label in each of the movies. Figure 5 shows the distribution of the *p*-values of the t-tests carried out on the individual movies. Results vary greatly across movies for all three features, but particularly for the eyes and mouth. Next, we analyzed the effects of additional movie data on between-group fixation differences. We randomly selected three movie clips from our 22 movies and performed a two-sample t-test on the average fixation times on each of the face labels between the ASD and TD groups in this movie set. This process was repeated for 10,000 permutations. Similar to the previous analysis, we examined the effect of incremental additions of data by repeating this process with random combinations of five, eight and 11 movies. Figure 6 features the distribution of *p*-values for eye, mouth, and nose fixations as the number of movie clips increases. As displayed in the histograms, the distributions become increasingly narrower as more movies are added. For mouth labels, the percentage of results showing significant differences between ASD and TD fixations decreases as the number of movies increases from one (9%) to 11 movies (0.98%), though we see no change in eye labels between one (0%) and 11 movies (0%). For nose labels, we observe an increase in the percentage of results showing significant differences between ASD and TD groups as the number of movies increases from one (53.9%) to 11 (100%) movies. This is further evidenced by permutation test results comparing differences in the effects of additional movie data on between group distributions per face label. Findings reveal significant differences between each of the distributions on both median and variance (*p*_median_ < 1×10^−4^; *p*_variance_ < 1×10^−4^ for all).

**Figure 5.**
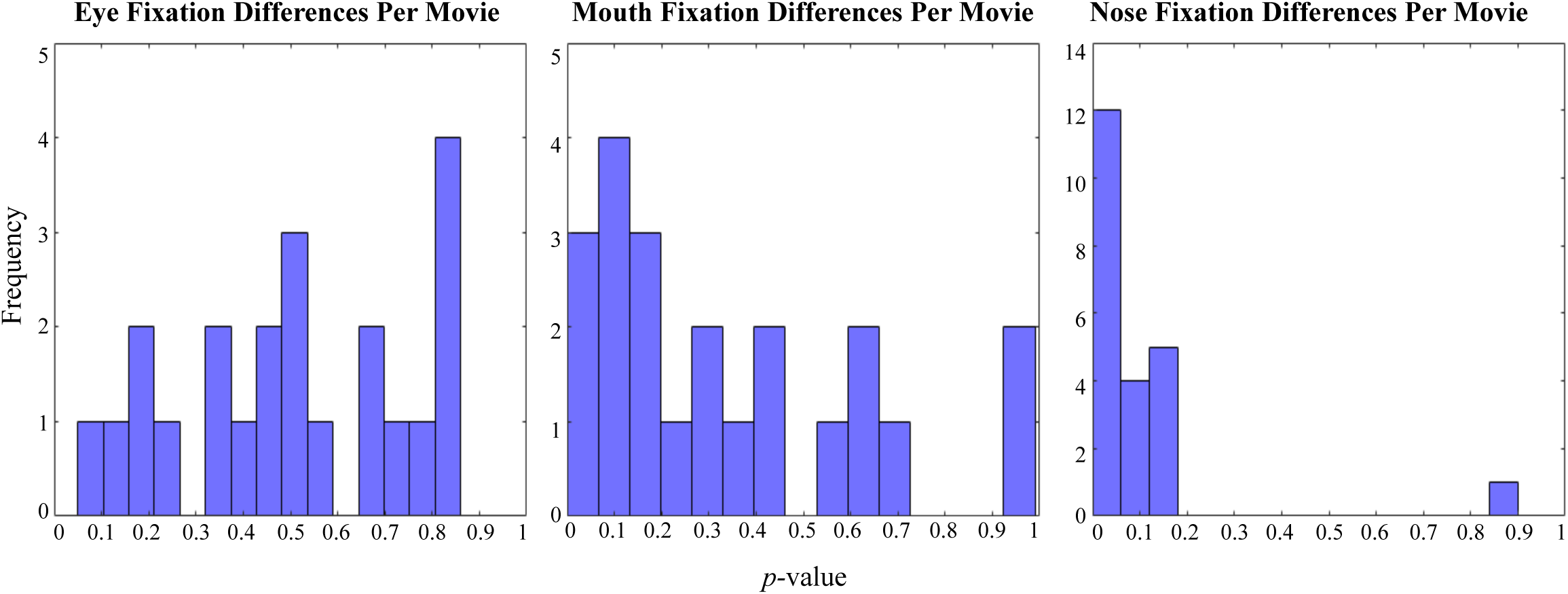
Distribution of the variability (t-test *p*-values) between TD individuals and individuals with ASD based on separate evaluation of each of the 22 movie clips. Results vary greatly across movies for all three features, but particularly for the eyes and mouth.

**Figure 6.**
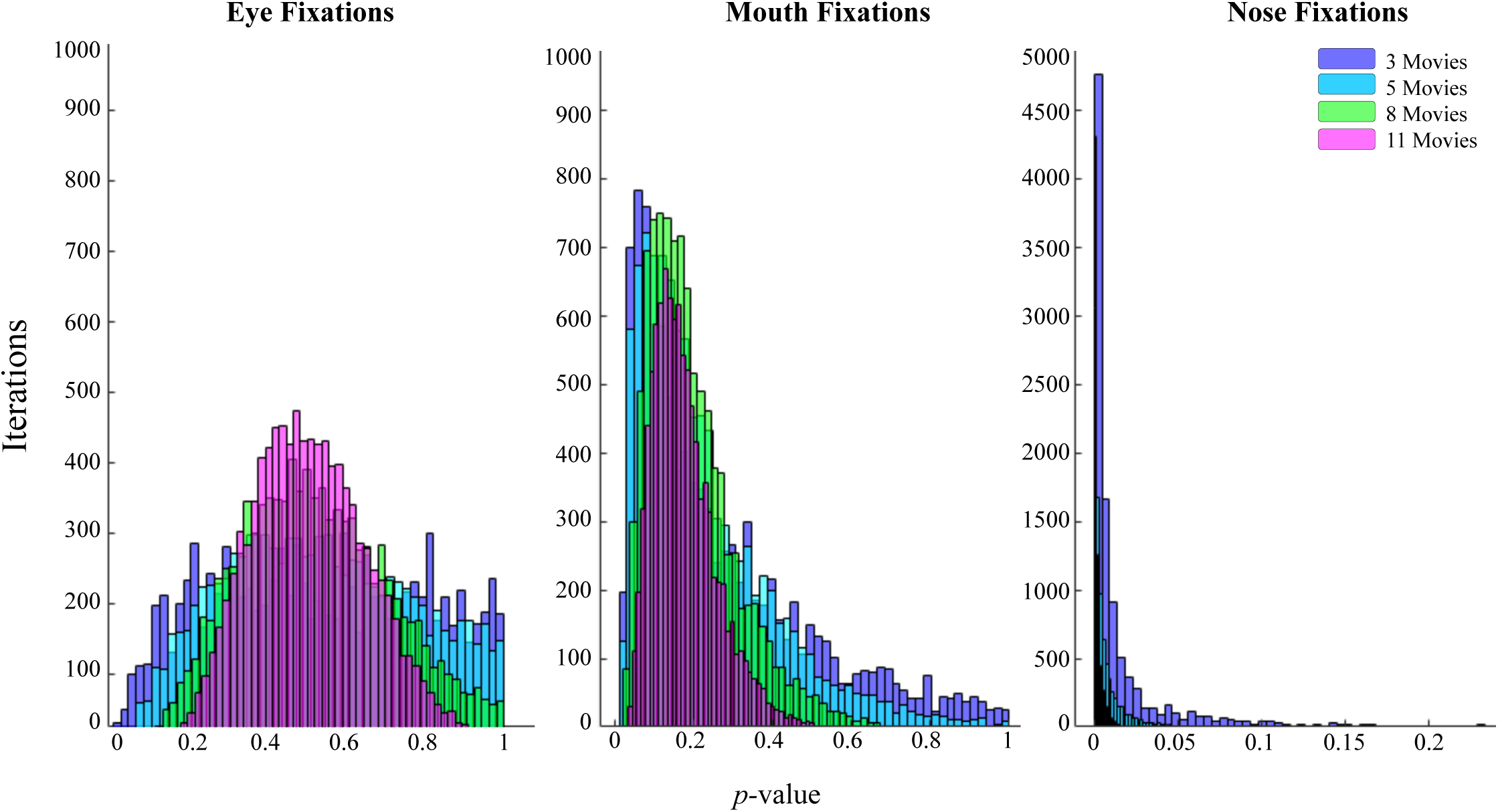
Consistency of fixations for TD individuals and those with ASD across movies when introducing additional movie data. Histograms reflect the distribution of t-test *p*-values of all permutations as the number of movie clips increases for eye, mouth, and nose fixations, respectively (see methods for details).

### Fixation of social interactions

Using 22 movie clips (308 seconds of stimuli) shown by the previous analysis to yield consistent and robust results, we sought to assess whether fixations of TD participants and those with ASD differ while viewing facial features during naturalistic dynamic interactions. Figure 7 displays a distribution of ASD and TD time spent fixating on each individual face label. With a two-by-three analysis of variance, we examined if fixation time is affected by diagnosis (ASD/TD) and individual core facial feature (eyes/nose/mouth). Main effect analyses revealed significant differences among face labels (*F* (1, 206) = 120.26, *p* = 4.49×10^−35^) and diagnosis (*F* (1, 206) = 5.22, *p* = .02). There was a statistically significant interaction between effects of diagnosis and face label on fixation time (*F* (1, 206) = 4.46, *p* =.01). TD participants attend more to the nose (*t* = 3.52; *p* = 7.73 × 10^−4^), however there was no difference between ASD and TD eye (*p* = .50) and mouth (*p* = .14) fixation times.

**Figure 7.**
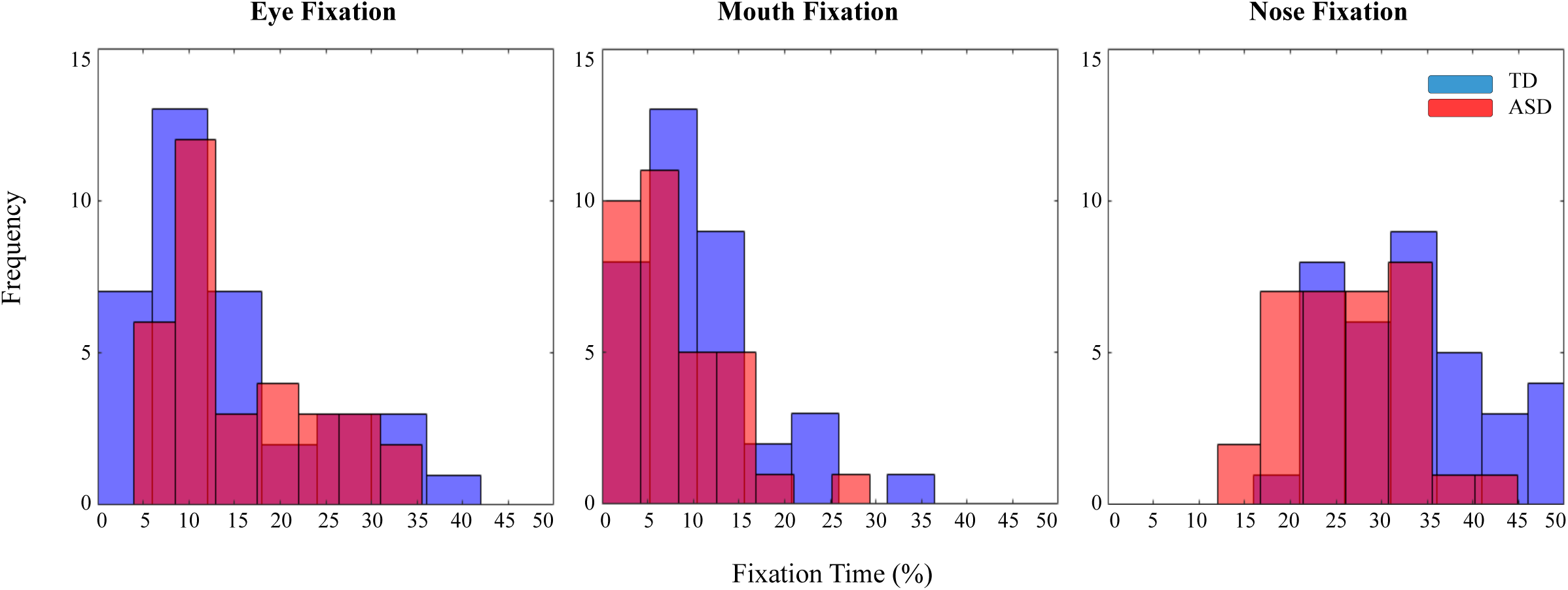
Distribution of proportion of time spent fixating on eye, mouth, and nose label (out of total face fixation time) for TD individuals and those with ASD using all 22 movie clips.

## Discussion

Our research investigates the stability of social fixations to complex and dynamic interactions in both TD individuals and those with ASD, while highlighting a machine learning approach to eye tracking paradigms. Overall, our findings demonstrate that fixation proportion to different facial features can be considered a stable trait when sufficient movie data are considered. As hypothesized, our ability to measure ASD and TD eye movement fixations gains consistency and robustness as we introduce incremental amounts of movie data to the analyses while using few movie clips yields unstable fixations to social stimuli. Based on a sufficient number of movie clips shown by our analysis to yield stable results (22 movie clips), we then sought to examine social fixation differences between TD individuals and those with ASD. Our findings reveal that individuals with ASD attend less to the center of the face (the nose region), and do not differ from TD individuals in eye and mouth fixation times. These results provide a feasible approach to portraying robust fixations of social viewing in individuals with ASD.

Eye tracking paradigms operate under the assumption that gaze fixations are a stable trait. However, previous literature prods at this assumption, showing that eye fixations vary depending on type of stimuli (static versus dynamic)^7^. Our results further drive at this question by revealing poor internal consistency when too few data are used. Although previous studies show individuals’ gaze consistency in static images, our stimuli feature rich content from several movies, and may reveal complexities that arise when using dynamic stimuli^15,19^. As seen by the relationship between single movie pairings, there is considerable individual variation from movie to movie (Figure 3a - c). Both diagnostic groups display this variability, though individuals with ASD display greater instability between individual movies as compared to TD counterparts across all face labels (Figure 3a - c). Nevertheless, our findings also depict the growing stability of fixation findings when introducing additional movie data for study. We see growing consistency within a participant to his own fixations as we increase the number of movies considered (Figure 4). Thus, we assert that with sufficient data our estimates of fixation proportions can be considered a stable trait in those with ASD and TD individuals. It is important to note that even with the addition of data, those with ASD still display greater variance in consistency than TD individuals (Figure a - c). This reflects that while fixations can be considered stable for those with ASD, more data is needed to see suitable stability in fixations among this group.

It is important to note that our data utilizes different, short movie clips. While using heterogenous movies can support the generalizability of fixation proportion as a stable individual trait, there are many other elements which could affect fixation proportion which we did not test for in this study. These data were all collected in a single session, and may not capture individual variation across days, though previous studies have shown some stability in this regard^19^.

Similarly, all the movies depict social interactions, and the task was a free viewing task. Different task context or very different movie content may also affect gaze patterns^7^. Future studies may seek to examine how much data would be necessary to robustly estimate individual fixation preferences when using a single, longer movie clip.

As hypothesized, examining the effect of movie data amount on the reliability of ASD and TD fixations differences revealed that results gained significant stability as we incrementally introduced clips for study. When using only a single short movie clip, findings widely vary in possible between group analyses results (Figure 5) and potentially yield both false-positive and false-negative significant group differences. We observe increased stability in the t-test results between groups as movies are added to the dataset being considered. Using 11 movie clips, the differences in mouth fixation times between TD and ASD participants were overwhelmingly non-significant (99%), whereas using only 1 movie clip yielded significant differences 9% of the time (Figure 6). In contrast, significant differences between ASD and TD nose fixation times were found for all possible movie set combinations when examining 11 movies but failed to reach significance when examining 46% of the single movies. As expected based on these results, using all 22 movies, group differences in nose fixation times were significant, but group differences in eye and mouth fixations were not.

While the stability of the between-group differences increases across all facial features with the addition of more data, there are clear differences in the distributions across labels. Distributions for eye and mouth fixation differences remain quite broad throughout, spanning both significant and insignificant results even with 5 movie clips. Distributions for nose fixation differences are much narrower, with all possible combinations of 5 movie clips yielding significant between-group differences (Figure 6). This likely indicates an interaction between the internal consistency of the individual data and the effect size of between-group differences.

As previously mentioned, there is wide and inconsistent debate on the extent to which individuals with ASD avoid eyes in favor of the mouth^2,3,4,5,6^. Our results emphasize the notable probability for misinterpretation of fixation data when using inadequate stimuli. The pliability of eye tracking findings raises a concern for the eye tracking field’s wide range of results; these results may provide greater commentary on stimuli dependent findings than on social processing in ASD.

By determining the amount of movie data shown to be sufficient for observing reliable differences, we then were able to reliably examine how individuals with ASD view elements of social interactions relative to TD peers. Our aberrant gaze findings demonstrate disrupted social viewing associated with ASD (Figure 7). Deviations in fixations are likely a reflection of ASD deficits in neural systems that modulate complex social behaviors, as evidenced by our previous research in which we link aberrant eye movements and atypical neural mechanisms in the “social brain”^20^. This is also supported by Avni and colleagues (2019), whose findings report reduced eye movement typicality in those with ASD, as well as a correlation between individual gaze idiosyncrasies and ASD severity^21^. Further, in this previous study, participants with ASD significantly vary in the overall typicality of eye movement scan paths compared to their TD counterparts. The present study elaborates on the typicality of ASD viewing fixations by featuring a specific finding in which individual behavior varies; our participants with ASD display significantly greater within-group variance in time spent fixating on the face, compared to TD individuals, as well as significantly reduced internal consistency in the viewing of the different facial features. Given well-established idiosyncrasies within the ASD population, this variance may reflect a behavioral manifestation of heterogeneous profiles within this disorder^22,23,24^.

Generally, the existing eye tracking literature substantiates findings of decreased ASD gaze fixations to socially relevant stimuli (i.e. facial features)^2^. Moreover, some findings place particular emphasis on differences in eye and mouth fixation times, and state that atypical viewing patterns of these features are characteristic and even predictive of autism^5^. However, we found no evidence to show that TD individuals and high-functioning individuals with ASD differ on either eye or mouth fixation time. Although individuals with ASD generally spend less time overall fixating on the face, it appears to be the nose which drives these facial region fixation differences; TD individuals fixate on this central facial feature significantly more than participants with ASD.

As evidenced by previous work, TD gaze allocation towards the nose may demonstrate several visual tendencies that are typical in normative populations. First, findings show that TD individuals initially fixate on the geometric center of the face (i.e. the eye-nose region) before exploring other features^25^. Rogers and colleagues (2018) report the existence of an “eye-mouth gaze continuum” in which TD individuals experiencing real-world interactions distribute their gaze in the area between eye and mouth regions, with variation in specific feature preference^26^. TD facial perception studies commonly exhibit this scan path when participants undergo face perception tasks^25,27^. Findings also show TD individuals display preferential attention to the area around the center of the nose during face recognition tasks^27^. In our study, the presence of nose fixations during movie watching may reflect TD individuals’ attempt to distinguish each scene’s various characters. This nose fixation is shown to provide a central point where the viewer’s periphery can take in information from the entire face^27^. This optimizes face perception and demonstrates the holistic nature of recognizing faces in TD individuals.

Nose-looking fixations appear typical and relevant to holistic face processing. Therefore, it is worth noting the diminished nose-looking behavior found in those with ASD within the present study. Based on what is known about the centrality of nose fixations, the lack thereof in those with ASD may suggest local processing with a bias towards local facial features. Prior ASD research not only reports evidence for local bias in visual perception, but also suggests that local processing tendencies in autism may contribute to the associated overall difficulty with integrating features to create a global representation^28,29^. Reduced nose-looking may reveal a developmental behavior that results from atypical social brain neural systems.

It should be noted that the constraints of manual stimuli segmentations (e.g. time-consuming, subject to human error) make it commonplace for studies to only include eye and mouth regions in facial feature coding. We expand on previous work by acknowledging the eye, mouth, and nose regions in our core facial feature analysis, which in turn revealed data-driven results that diverge from findings of exclusive eye- and mouth-directed analyses^7,9^.

The current study makes methodological advances in the use of eye tracking as a tool in detecting ASD gaze abnormalities. Previous studies primarily use manual segmentation techniques which severely limit our ability to reproduce methods and compare findings across studies. Other investigators seeking to use eye movements as biomarkers in ASD research acknowledge the challenge of aggregating findings and have thus called for objective and quantifiable outcome measures^14^. Additionally, the constraints of manual stimuli segmentation lead researchers to reduce the amount of eye tracking stimuli or reuse already-segmented clips, however our results demonstrate the pitfalls of insufficient data on fixation results. The present study’s machine learning algorithm fulfills the need for a quantifiable and data-driven approach and optimizes the use of ample and diverse stimuli, while eliminating the typical restrictions of manual techniques. We encourage future studies to adopt similar automatic stimuli segmentation techniques to enable the use of the large amounts of stimuli needed to reliably test hypothesis about social gaze processing in populations.

Overall, we assessed the stability of ASD social processing gaze fixations using a data-driven machine learning approach to dynamic eye tracking stimuli segmentation. Our ability to estimate fixations of TD individuals and those with ASD gains robustness and consistency as we introduce additional amounts of movie data for study, while few movie clips yield unstable fixations to social stimuli. Based on a sufficient amount of movie data, we conclude that individuals with ASD do not attend to social interactions the same way as their typically developing peers. Individuals with ASD do not differ on eye or mouth fixations compared to TD peers, rather they exhibit less centralized fixations when viewing the face. Naturalistic stimuli, paired with a machine learning approach for reliable segmentations, can advance the technical capability of eye movement analysis.

## Methods

### Participants

Fifty high-functioning males with ASD and 36 TD male participants were recruited for this study at the National Institute of Mental Health (ClinicalTrials.gov: NCT01031407). All participants with ASD met the cutoff for the category designated as “broad autism spectrum disorders” according to the criteria established by the National Institute of Child Health and Human Development/ National Institute on Deafness and Other Communication Disorders Collaborative Programs for Excellence in Autism^30^. Seventeen participants with ASD were omitted from this analysis due to incomplete testing data (n = 5), poor quality eye tracking data (n = 4), did not meet autism diagnosis (n = 3), scheduling conflicts (n = 2), did not meet IQ cut off (Full Scale IQ > 70; n = 1), conflicting medical conditions (n = 1), and loss to follow up (n = 1). TD participants were selected to create an age- and IQ-equated match for each participant with ASD. TD participants and participants with ASD did not differ on age, IQ, race, or ethnicity (Table 1).

**Table 1.** Demographics chart. There are no statistically significant differences between participants with ASD and TD participants across age, IQ, race, or ethnicity.

|  | ASD (n = 33) | TD (n = 36) | Total (n = 69) |
| --- | --- | --- | --- |
| Age, mean (SD) | 20.25 (4.02) | 20.83 (3.82) | 20.74 (4.0) |
| IQ, mean (SD) | 107.81 (13.75) | 108.82 (12.98) | 108.23 (13.11) |
| Race, n (%) |  |  |  |
| White | 21 (63.6) | 20 (55.5) | 41 (59.4) |
| Black | 2 (6.1) | 6 (16.7) | 8 (11.6) |
| Asian | 1 (3.0) | 3 (8.3) | 4 (5.8) |
| Biracial | 3 (9.1) | 4 (11.1) | 7 (10.15) |
| Other | 1 (3.0) | 1 (2.8) | 2 (2.9) |
| Unknown | 5 (15.2) | 2 (5.6) | 7 (10.15) |
| Ethnicity, n (%) |  |  |  |
| Hispanic/ Latino | 6 (18.2) | 5 (13.9) | 11 (15.9) |
| Not Hispanic/ Latino | 22 (66.7) | 30 (83.3) | 52 (75.4) |
| Unknown | 5 (15.1) | 1 (2.8) | 6 (8.7) |

### Procedure

Participants’ heads were stabilized using a forehead and chin rest, and eye gaze calibrations were performed on the right eye at the beginning of the experiment. Participants engaged in an 8-minute free-viewing paradigm. There were no explicit instructions other than to watch the presented movies. They viewed 24 movie clips (14 seconds in duration) depicting social interactions in which two or more characters engage in conversation. Movie clips consisted of the following Hollywood movies: The Blind Side (6 clips), The Goonies (4 clips), How To Lose a Guy in Ten Days (4 clips), The Italian Job (5 clips), and The NeverEnding Story (5 clips). Movies were viewed full screen on a digital monitor with a 1920 × 1080 resolution, with a screen size of 20.5 × 12 inches. Eye movements were recorded by the Eyelink 1000 Plus, sampled at 1000 Hz. A grey screen appeared for 6 seconds between presentations of the clips. A fixation cross appeared in the center of the grey screen to reset fixations to the center before presenting the successive clip.

For our analyses, we excluded two movie clips from The NeverEnding Story for displaying a highly disproportionate ratio of face-to-background pixels or scene darkness that altered segmentations. Final analyses included 22 movies (7,410 frames).

### Image Segmentation

We trained an ANN to predict segmentations of each pixel for each frame for each movie. We used the Pascal-Parts dataset to train a Bayesian SegNet with concrete dropout to make a predicted segmentation for a given movie frame (Figure 1)^31,32,33^. When applying the ANN to new movie frames, 10 concrete dropout Monte-Carlo samples were used to produce predicted segmentation labels and uncertainty. Figure 2 displays a comparison of segmented stimuli to the original frame. The code we used is publicly available at https://github.com/nih-fmrif/MLT_Body_Part_Segmentation, and further details about the ANN, see McClure, Reimann, Ramot, & Pereira (In preparation)^34^.

**Figure 1.**
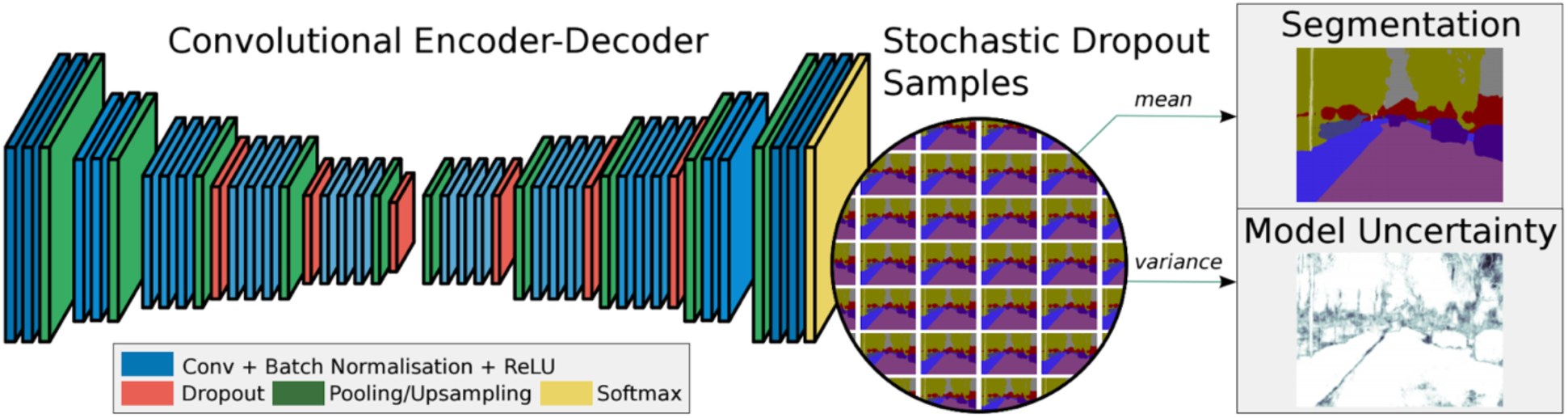
An illustration of a Bayesian SegNet artificial neural network (ANN) and the sampling process used to generate a predicted segmentation and uncertainty (from Kendall et al. (2015).

**Figure 2.**
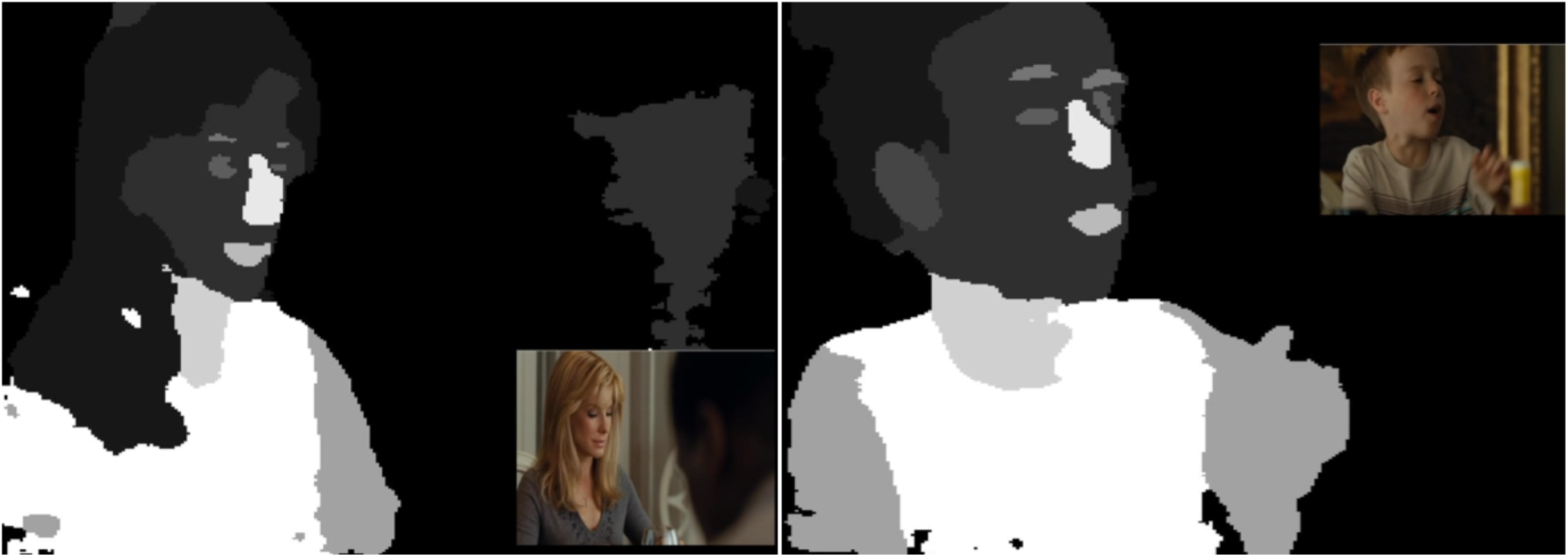
Artificial Neural Network (ANN) identified and segmented each frame of the dynamic stimuli. Examples comparing two original movie frames to the segmented image in black and white. Different shades of white/grey represent our varying labels generated by the ANN. Black segments represent no labels given by the ANN, which in our analysis was then labeled as background.

The ANN segmented images into 11body part labels: hair, head, ear, eye, eyebrow, leg, arm, mouth, neck, nose, and torso. Additionally, we created a 12th category for each pixel that the ANN did not place into one of these 11 labels. This label was treated as the background label and contained all other frame features such as objects, landscapes, and noise that were not were not associated with the 11 other labels. The performance of the ANN was tested on the test set portion of the Pascal-Parts dataset. The average Dice score across body part labels was 0.55, and 0.95 for background^31,34^.

### Eye tracking processing

Eye movement data was extracted for each separate movie clip, removing the first and last 500 milliseconds of each clip. Non-fixation data (e.g. blinks, missing or offscreen fixations) were ignored. Data were despiked and sampled down to the frame rate at which the clips were presented (29.97 frames per second). Each pixel received one of the aforementioned 12 labels based on ANN output. To categorize each fixation as one of these labels, we examined the algorithm’s label predictions within the 15-pixel radius surrounding the primary fixation point. After which, the most frequently occurring pixel label was selected with a bias towards smaller features; for example, if the pixels within a 15-pixel radius from a particular fixation included both ‘eye’ and ‘head’ labels, that fixation would be labeled as ‘eye’.

### Data Analysis

For the purposes of this analysis, we examined social fixations to the core facial features including eyes, nose, and mouth. This analysis investigated the effects of varying amounts of movie data on the consistency of individual fixations to these core features as well as between-group differences. As a basis for evaluating consistency of fixations across movie clips with varying content, we normalized the gaze data for each movie clip. For each participant and each movie clip, we calculated the proportion of time spent looking at each individual face label (eyes, nose, mouth) divided by total face fixation time (time spent fixating on eyes, nose and mouth labels together). For each participant, movie clip, and facial feature, we then calculated the distance from the average fixation time of all other participants in their respective groups.

This was normalized by total time spent looking at the face, as described above. This normalization serves to account for differences in raw fixation time on the different facial features; these differences may arise from movie-specific variability (e.g. number of feature pixels per frames, action or speech content) and draw attention to or from the different features. The resulting values for each participant (henceforth referred to as fixation proportion represent the proportion of looking time they allocate to each of the facial features out of the time they spend looking at the face in general for that particular movie clip, compared to all other participants in their group. These values were then used to evaluate the internal consistency of fixation time on each of the facial features across movie clips. This was done by correlating the fixation proportion for participants across different movie pairs / movie sets, for all possible combination of single movie pairs, and for 10,000 randomly selected movie sets of three, five, eight, and 11 movies.

### Statistical analysis

To evaluate differences in the distributions between proportion of time spent looking at core features, we performed permutation-based statistical tests using movie sets consisting of one, three, five, eight, and 11 different movie clips; these analyses were repeated for each face label (eyes, nose, and mouth). To test whether the ASD and TD correlation coefficient distributions significantly differ from each other, we first calculated the TD group’s median fixation proportion subtracted by the ASD group’s median fixation proportion (henceforth known as real median differences), as well as the TD fixation proportion variance subtracted by ASD fixation proportion variance (henceforth known as real variance differences). We then generated two sets of 10,000 randomly selected fixation proportions from combined ASD and TD eye tracking data, by randomly permuting the TD and ASD labels. From this permuted dataset, we calculated the first dataset’s median fixation proportion subtracted by the second dataset’s median fixation proportion (henceforth known as permuted median differences), as well as the first dataset’s fixation proportion variance subtracted by the second dataset’s fixation proportion variance (henceforth known as permuted variance differences). This process was repeated 10,000 times. For each iteration of permuted differences, we calculated the proportion of permuted median differences greater than real median differences, as well as the proportion of permuted variance differences greater than real variance differences. This resulting number represents a two-tailed *p*-value.

For analysis of within-group fixation stability, we randomly selected two sets of three non-overlapping movie combinations, totaling 42 seconds of stimuli. Similar to the aforementioned analysis, we calculated the correlations of the within-group internal consistency of the fixation proportions across these two sets of movies; this was done for TD participants and participants with ASD separately. This process was repeated for 10,000 permutations, with different sets of three movies selected for each permutation; to assess incremental additions of data, the process was repeated with two sets of movies with random combinations of five (70 seconds), eight (112 seconds), and 11 (154 seconds) movies. Then, we sought to evaluate if these correlation coefficient distributions for each varying level of movie data (three, five, eight, and 11 movies) significantly differ from each other. Differences between each of the movie data distributions refers to the following comparisons: three versus five movies, three verses eight movies, three versus 11 movies, five versus eight movies, five versus 11 movies, and eight versus 11 movies. For each of these pairwise combinations, we calculated the real median differences between the first movie level and the second movie level, as well as the real variance differences between the first movie level and the second movie level. We then generated two sets of 10,000 randomly selected fixation proportions from combined movie level eye tracking data. From this permuted dataset, we calculated the permuted median differences between the first dataset and the second dataset, as well as the permuted variance differences between the first dataset and the second dataset. As before, this process was repeated 10,000 times as we calculated the proportion of permuted median differences greater than real median differences, as well as the proportion of permuted variance differences greater than real variance differences. This resulting number represents a two-tailed *p*-value.

For analysis of between-group fixation stability, we randomly selected two sets of non-overlapping movie combinations; this was done on sets of three, five, eight, and 11 movies as described above. For 10,000 permutations of these randomly selected sets, we calculated the correlations of the within-group internal consistency of the fixation proportions across these two sets of movies; this was done for TD participants and participants with ASD separately. Then, we sought to evaluate if these ASD and TD correlation coefficient distributions significantly differ from each other across akin levels of movie data. First, we calculated the real median differences between TD and ASD data, as well as the real variance differences between TD and ASD data. As before, we generated two sets of 10,000 randomly selected fixation proportions from combined movie level eye tracking data. From this permuted dataset, we calculated the permuted median differences between TD and ASD data, as well as the permuted variance differences between TD and ASD data. As before, this process was repeated 10,000 times as we calculated the proportion of permuted median differences greater than real median differences, as well as the proportion of permuted variance differences greater than real variance differences. This resulting number represents a two-tailed *p*-value.

## Acknowledgments

This work was supported by the National Institute of Mental Health (ClinicalTrials.gov: NCT01031407). This material is based upon work supported by the National Science Foundation Graduate Research Fellowship under Grant No. DGE-1646737 and under Grant Nos. DGE-1650604 and DGE-2034835. These organizations did not have a role in the conceptualization, design, data collection, analysis, decision to publish, or preparation of the manuscript.

## References

1. Freeth, M., Chapman, P., Ropar, D., & Mitchell, P. (2010). Do gaze cues in complex scenes capture and direct the attention of high functioning adolescents with ASD? Evidence from eye-tracking. J. Autism Dev. Disord. 40, 534–547 (2010).

2. Yi, L., Fan, Y., Quinn, P.C., Feng, C., Huang, D., et al. Abnormality in face scanning by children with autism spectrum disorder is limited to the eye region: evidence from multi-method analyses of eye tracking data. J. Vis. 13, 5 (2013).

3. Gillespie-Smith, K., Riby, D.M., Hancock, P.J., & Doherty-Sneddon, G. Children with autism spectrum disorder (ASD) attend typically to faces and objects presented within their picture communication systems. J. Intellect. Disabil. Res. 58, 459–470.(2014).

4. Snow, J., Ingeholm, J.E., Levy, I.F., Caravella, R.A., Case, L.K., et al. Impaired visual scanning and memory for faces in high-functioning autism spectrum disorders: it’s not just the eyes. J. Int. Neuropsychol. Soc. 17, 1021–9 (2011).

5. Jones, W., Carr, K., & Klin, A. Absence of preferential looking to the eyes of approaching adults predicts level of social disability in 2-year-old toddlers with autism spectrum disorder. Arch. Gen. Psychiatry. 65, 946–54 (2008).

6. Asberg Johnels, J., Gillberg, C., Falck-Ytter, T., & Miniscalco, C. Face-viewing patterns in young children with autism spectrum disorders: speaking up for the role of language comprehension. J. Speech Lang. Hear. R. 57, 2246–2252 (2014).

7. Speer, L.L., Cook, A.E., McMahon, W.M., & Clark, E. Face processing in children with autism: Effects of stimulus contents and type. Autism. 11, 265–277 (2007).

8. van der Geest, J.N., Kemner, C., Verbaten, M.N., & van Engeland, H. Gaze behavior of children with pervasive developmental disorder toward human faces: a fixation time study. J. Child Psychol. Psychiatry. 43, 669–78 (2002).

9. Rice, K., Moriuchi, J.M., Jones, W., & Klin, A. Parsing heterogeneity in autism spectrum disorders: visual scanning of dynamic social scenes in school-aged children. J. Am. Acad. Child Adolesc. Psychiatry. 51, 238–248 (2012).

10. Wang, W., Liu, C., & Zhao, D. How much data are enough? A statistical approach with case study on longitudinal driving behavior. IEEE Trans. Intell. Veh. 99, (2017).

11. Heyman, R.E., Chaudhry, B.R., Treboux, D., Crowell, J., Lord, C. et al. How much observational data is enough? An empirical test using marital interaction coding. Behav. Ther. 32, 107–122 (2001).

12. Splinter, K.D., Turner, I.L., & Davidson, M.A. How much data is enough? The importance of morphological sampling interval and duration for calibration of empirical shoreline models. Coast. Eng. 77, 14–27 (2013).

13. Wortley, A.H., Rudall, P.J., Harris, D.J., & Scotland, R.W. How much data are needed to resolve a difficult phylogeny? Case study in lamiales. Syst. Biol. 54, 697–709 (2005).

14. Bradshaw, J., Shic, F., Holden, A.N., Horowitz, E.J., Barrett, A.C., et al. The use of eye tracking as a biomarker of treatment outcome in a pilot randomized clinical trial for young children with autism. Autism Res. 12, (2019).

15. Arizpe, J., Walsh, V., Yovel, G., & Baker, C.I. The categories, frequencies, and stability of idiosyncratic eye-movement patterns to faces. Vis. Res. 141, 191–203 (2017).

16. Peterson, M.F. & Eckstein, M.P. Individual differences in eye movements during face identification reflect observer-specific optimal points of fixation Psychol. Sci. 24, 1216–1225 (2013).

17. Poynter, Barber Inman, & Wiggins. Individuals exhibit idiosyncratic eye-movement behavior profiles across tasks. Vis. Res. 89, 32–38 (2013).

18. Castelhano, M., & Henderson, J. Stable individual differences across images in human saccadic eye movements. Can. J. Exp. Psychol. 62, 1–14 (2008).

19. Mehoudar, E., Arizpe, J., Baker, C.I., & Yovel, G. Faces in the eye of the beholder: Unique and stable eye scanning patterns of individual observers. J. Vis. 14, 1–11 (2014).

20. Ramot, M., Walsh, C., Reimann, G.E., & Martin, A. Distinct neural mechanisms of social orienting and mentalizing revealed by independent measures of neural and eye movement typicality. Commun. Biol. 3, (2020).

21. Avni, I., Meiri, G., Bar-Sinai, A., Reboh, D, Manelis, L., et al. Children with autism observe social interactions in an idiosyncratic manner. Autism Res. 13, 935–946 (2020).

22. Hasson, U., Avidan, G., Gelbard, H., Vallines, I., Harel, M., et al. Shared and idiosyncratic cortical activation patterns in autism revealed under continuous real-life viewing conditions. Autism Res. 2, (2009).

23. Hahamy, A., Behrmann, M., & Malach, R. The idiosyncratic brain: Distortions of spontaneous connectivity patterns in autism spectrum disorder. Nat. Neurosci. 18, 302–309 (2015).

24. Byrge, L., Dubois, J., Tyszka, J.M., Adolphs, R., & Kennedy, D.P. Idiosyncratic brain activation patterns are associated with poor social comprehension in autism. J. Neurosci. 35, 5837–5850 (2015).

25. Bindemann, M., Scheepers, C., & Burton, A.M. Viewpoint and center of gravity affect eye movements to human faces. J. Vis. 9, (2009).

26. Rogers, S.L., Speelman, C.P., Guidetti, O., & Longmuir, M. Using dual eye tracking to uncover personal gaze patterns during social interaction. Sci. Rep. 8, 4271 (2018).

27. Hsaio, J.H. & Cottrell, G. Two fixations suffice in face recognition. Psychol. Sci. 19, 998–1006 (2008).

28. Nayar, K., Voyles, A.C., Kiorpes, L., & Di Martino, A. Global and local visual processing in autism: An objective assessment approach. Autism Res. 10, 1392–1404 (2017).

29. Shah, P., Bird, G., & Cook, R. Face processing in autism: Reduced integration of cross-feature dynamics. Cortex. 75, 113–119 (2016).

30. Lainhart, J.E., Bigler, E.D., Bocian, M., Coon, H., Dinh, E. et al. Head circumference and height in autism: a study by the Collaborative Program of Excellence in Autism. Am. J. Med. Genet. Part A. 140, 2257–2274 (2006).

31. Everingham, M., Van Gool, L., Williams, C. K., Winn, J., & Zisserman, A. The Pascal visual object classes (voc) challenge. Int. J. Comput. Vis. 88, 303–338 (2010).

32. Kendall, A., Badrinarayanan, V., & Cipolla, R. Bayesian segnet: Model uncertainty in deep convolutional encoder-decoder architectures for scene understanding. arXiv (2015).

33. Gal, Y., Hron, J., & Kendall, A. Concrete dropout. Adv. Neural Inf. Process. Syst. 3581–3590 (2017).

34. McClure, P., Reimann, G.E., Ramot, M., & Pereria, F. A deep neural network tool for automatic segmentation of human body parts in natural scenes. arXiv. arXiv:2009.09900

